# Single-molecule genome assembly of the Basket *Willow, Salix viminalis*, reveals earliest stages of sex chromosome expansion

**DOI:** 10.1101/589804

**Authors:** Pedro Almeida, Estelle Proux-Wera, Allison Churcher, Lucile Soler, Jacques Dainat, Pascal Pucholt, Jessica Nordlund, Tom Martin, Ann Christine Rönnberg-Wästljung, Björn Nystedt, Sofia Berlin, Judith E. Mank

## Abstract

Sex chromosomes have evolved independently multiple times in eukaryotes and are therefore considered a prime example of convergent genome evolution. Sex chromosomes are known to emerge after recombination is halted between a homologous pair of chromosomes and this leads to a range of non-adaptive modifications causing the gradual degeneration and gene loss on the sex-limited chromosome. However, because studies on sex chromosomes have primarily focused on old and highly differentiated sex chromosomes, the causes of recombination suppression and the pace at which degeneration subsequently occurs remain unclear. Here, we use long- and short-read single molecule sequencing approaches to assemble and annotate a draft genome of the basket willow, *Salix viminalis*, a species with a female heterogametic system at the earliest stages of sex chromosome emergence. Our single-molecule approach allowed us to phase the emerging Z and W haplotypes in a female, and we detected very low levels of Z/W divergence, largely the result of the accumulation of single nucleotide polymorphisms in the non-recombining region. Linked-read sequencing of the same female and an additional male (ZZ) revealed the presence of two evolutionary strata supported by both divergence between the Z and W haplotypes and by haplotype phylogenetic trees. Gene order is still largely conserved between the Z and W homologs, although a few genes present on the Z have already been lost from the W. Furthermore, we use multiple lines of evidence to test for inversions, which have long been assumed to halt recombination between the sex chromosomes. Our data suggest that selection against recombination is a more gradual process at the earliest stages of sex chromosome formation than would be expected from an inversion. Our results present a cohesive understanding of the earliest genomic consequences of recombination suppression as well as valuable insights into the initial stages of sex chromosome formation.

## Introduction

Sex chromosomes, genomic regions associated with either males or females, have evolved independently many times in the eukaryotes [1, 2]. Sex chromosomes come in two general forms in organisms where sex is expressed in the diploid phase of the life cycle. X-Y sex chromosomes form where the Y chromosome is associated with males (male heterogamety), and Z-W sex chromosomes form where the W is associated with females (female heterogamety). Both these sex chromosome types emerge after recombination is halted between a homologous pair of chromosomes [3, 4], which allows the X and Y or Z and W chromosomes to diverge from each other. Studies in systems with unrelated and highly diverged sex chromosomes have revealed many shared genomic properties across a broad array of taxa [1, 2, 5], and sex chromosomes therefore represent an important example of convergent genome evolution.

In addition to promoting the sex chromosomes to diverge from one another, recombination arrest in the sex determining region (SDR) leads to a range of non-adaptive consequences for the sex-limited Y or W chromosome. These include the build-up of deleterious variation and repetitive elements, as well as loss of gene activity [6–8]. Due to the longstanding focus on systems with highly divergent sex chromosomes, the speed and order at which these processes occur after recombination suppression remain largely unclear.

Additionally, over evolutionary time, the non-recombining region can expand, resulting in strata, or regions with differing levels of divergence between the X and Y or Z and W chromosomes [9–13]. Expansion of the non-recombining region and the emergence of new strata may occur gradually, in which case we might expect only partial recombination suppression in the youngest stratum, in conjunction with substantial heterogeneity in X-Y or Z-W divergence [14–17]. Alternatively, some have suggested that strata form instantaneously, via large-scale inversions on the Y or W chromosome [18], which prevent recombination between the sex chromosomes along the entirely of the reversed region.

The answers to these questions have important implications beyond sex chromosomes. Halting recombination permanently links co-adapted gene complexes [19–22], also referred to as supergenes. Y and W chromosomes are thought to represent sex-specific supergenes, linking loci with sex-benefit alleles to the sex determining locus [23–26]. Supergenes have resurfaced recently as a major potential adaptive mechanism [27–31], and in so doing have implicated recombination suppression as a crucial component of complex phenotypic adaptation.

Sex chromosomes are therefore a powerful system to understand the evolutionary consequences of recombination suppression and supergene formation. Furthermore, detailed studies of nascent sex chromosomes are critical if we want to understand the initial causes of recombination suppression, as well as the order and rate of the evolutionary processes that follow it. For example, recent studies of young sex chromosome systems have revealed substantial intra-specific variation in the degree of recombination suppression across populations [32–35], suggesting a dynamism not normally observed in older, more diverged sex chromosome systems.

Plants in particular are useful in the study of the earliest stages of sex chromosome formation, as many plant sex chromosomes emerged only very recently in evolutionary time [36–39]. Recent studies based on next-generation sequencing of plant sex chromosomes have shown important patterns in the earliest stages of sex determination [40–45]. Studies on plant sex chromosomes have also revealed the importance of haploid selection in maintaining gene activity in the non-recombining region [15, 46] in the face of rapid loss of gene expression following recombination suppression [8, 47].

Recent work in *Salix viminalis*, the basket willow, has revealed the presence of nascent Z-W sex chromosomes, with a highly restricted SDR [48, 49]. The sex chromosomes of *Salix* have evolved independently from the X-Y sex chromosome system in the sister genus *Populus* [48, 50], which also exhibits very low levels of divergence [37]. The Salicaceae family, which includes willows and poplars, therefore presents a powerful system for studying the earliest stages of sex chromosome formation. Here, we use long- and short-read single-molecule sequencing (PacBio and 10x Genomics Chromium linked reads approaches) in *S. viminalis* to assemble a female reference genome. Importantly, our approach allowed us to obtain phased male and female haplotypes using large, continuous haplotype scaffolds. This allows us to transcend the current limitations of short-read next-generation sequencing, which hinder the assembly of repetitive regions, common in SDRs, as well as complicate accurate phasing. Our results shed unprecedented detail on the earliest stages of sex chromosome formation, and reveal that the initial stages of recombination suppression are incomplete, as would be expected from gradual selection against recombination rather than a single large-scale inversion.

## Results & Discussion

### Assembly and annotation of the Basket Willow reference genome

In order to gain a better understanding on the evolution and genomic architecture of the recently formed sex chromosomes in *Salix viminalis* we sequenced and assembled the complete genome of a single diploid heterogametic female (ZW) which was previously part of a large association mapping population [51]. To this end, we used a combination of long- and short-read single-molecule sequencing strategies and generated ∼19 Gb of Pacific Biosciences (PacBio) long reads in a female and ∼58 Gb of 10x Genomics linked-reads in the same female and a male (Table S1). The full assembled genome has ∼357 Mb of sequence spanning 2,372 scaffolds above 1 kb in length, an N50 value of ∼1.3 Mb and with 92% of the genome in scaffolds longer than 50 kb. With this estimated genome size, our sequencing constitutes >50X PacBio and >160X 10x Genomics coverage for autosomes, and >25X and >40X coverage of the W chromosome accounting for the hemizygous nature of the female-limited region.

Assembly quality, as assessed by whole-genome DNA and transcriptome short-read mapping, suggests a high completeness and contiguity with ∼98% and ∼84% of the reads respectively aligned to the assembled sequence (Table S2). Importantly, we obtained a high proportion of properly paired reads (Table S2). An initial assessment also identified more than 94% of complete core Embryophyta genes in the assembly (Table S2). We also mapped 1987 Genotype by Sequencing (GBS) [48, 52] markers in order to verify their presence and order. Consequently, our reference genome of the basket willow *S. viminalis* is essentially complete and properly assembled. Given the difficulties inherent in assembling an ancient polyploid genome such as *S. viminalis* [53], the relative completeness of our assembly reveals the benefits of incorporating single-molecule and long-read sequencing.

Annotation of the basket willow genome followed an in-house pipeline based on maker v3.00.0 [54] that combined transcriptome data [49, 55], reference proteins and *ab-initio* predictors. We identified 36,490 gene models, with 28,212 (77.3%) of them having functional annotation, and predicted 3,469 ncRNA and 1,139 tRNA (Table S3). Finally, we also identified several families of repetitive elements which together represent ∼35% of the assembly. The basket willow genome is publicly available for the community through the PopGenIE Integrative Explorer (http://popgenie.org) [56].

### Delimitation of the SDR in the female assembly

Differences between male and female genomes in read depth or single nucleotide polymorphim (SNP) density can be used to identify different forms of sex chromosome divergence [12, 57]. In nascent sex chromosome systems, this method is particularly useful when combined with genetic mapping studies of gender (in plants) or sex (in animals) [35, 49]. These methods are based on the different patterns of divergence and dosage between males and females on the sex chromosomes. In female heterogametic systems, W-specific reads are present only in females, resulting in higher female read coverage for W scaffolds. Conversely, as the W degrades, we expect a greater male read depth for the corresponding region of the Z chromosome, as females retain only one copy of the Z. Additionally, in the earliest stages of recombination suppression, we expect W-regions to retain significant similarity to the Z chromosome, and therefore females may show similar read depth for these regions as males. However, once recombination is halted, the W is expected to accumulate polymorphisms that are not shared with the Z, and so we might expect a greater density of SNPs in females compared to males in these regions even before significant divergence lowers mapping efficiency.

In order to assess these two levels of sex chromosome divergence, we mapped male and female short-read DNA-seq data (∼69X and ∼66X average coverage for females and males respectively) to our female assembly. Because we assembled the genome of a heterogametic Z-W female, and given the relatively high levels of heterozygosity across the genome (∼0.5% or 1 SNP per 200 bp), we expect a limited proportion of divergent regions in the genome, including Z and W haplotypes, to assemble separately in different scaffolds. As this would likely bias our SNP density estimates, where regions with elevated numbers of polymorphisms would appear to be homozygous, we first constructed a non-redundant assembly by removing smaller scaffolds that showed strong evidence of sequence overlap with longer scaffolds. We then aligned our non-redundant scaffolds to the *Populus trichocarpa* genome [58], revealing broad synteny as expected between these sister genera (Fig. S1, Fig. 1A). In total, we anchored ∼272 Mb (76.4% of the full assembly) to *P. trichocarpa* chromosomes.

**Figure 1.**
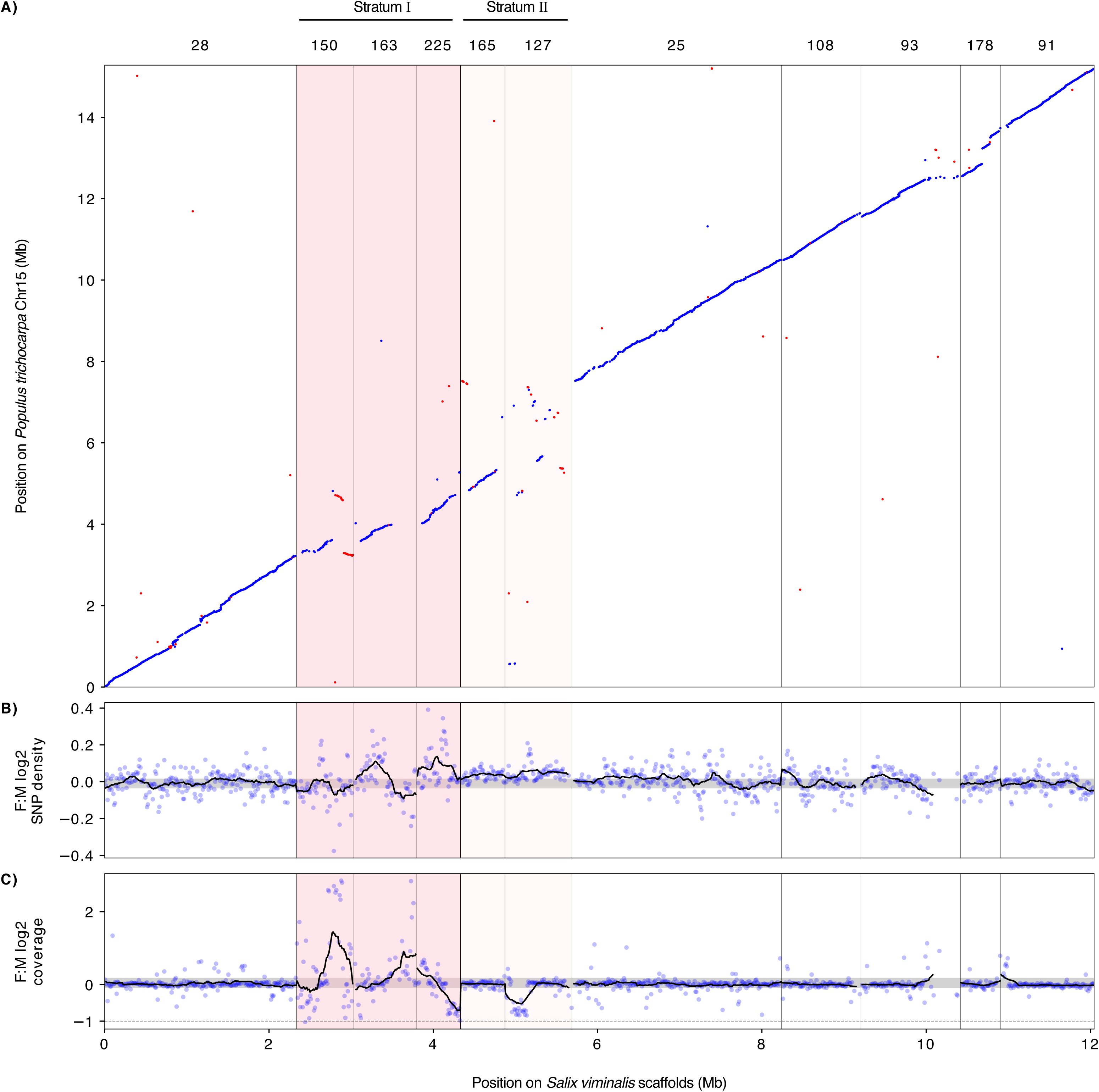
Identification of two evolutionary strata in the sex determination region (SDR) of the basket willow *S. viminalis*. Chromosome positions for *P. trichocarpa* and *S. viminalis* are shown in Mb with the *S. viminalis* scaffold names shown on the top. The two identified strata are shown with different hues of pink and labelled above the plot. **A)** Anchoring of *S. viminalis* scaffolds to the autosomal chromosome 15 of *P. trichocarpa*. Forward alignments are drawn in blue and reverse alignments are drawn in red. **B)** Log_2_ differences of normalized SNP density between *S. viminalis* females and males in non-overlapping windows of 10 kb. A moving average of 25 windows is shown in the black line. The grey shaded area corresponds to the bootstrap 95% confidence interval of the autosomal data. **C)** Log_2_ differences of normalized read coverage between females and males in non-overlapping windows of 10 kb. Moving average and bootstrap statistics are as in B). Values close to -1 indicate twice the coverage in males in comparison with female, thus potentially Z-linked.

We previously identified chromosome 15 as the sex chromosome [48, 49] and mapped the extent of the SDR on this chromosome (highlighted in pink, Fig. 1). Our results show that the five scaffolds within the SDR show significant deviations of both female:male SNP density, indicative of the build-up of female-specific SNPs on the W, and/or female:male read coverage differences, indicating regions of significant divergence between the Z and W chromosomes (Fig. 1). It is important to note that because *S. viminalis* exhibits only a limited divergence between the Z and W, and our long-read assembly was based on a female sample, the assembly of the sex chromosome regions likely represents Z-W chimeras. This chimerism is evident in scaffolds 150 and 163, which both show a region of similar coverage in males and females and a region of strong female-bias that likely represents W-specific genetic material (Fig. 1). These scaffolds, in addition to scaffold 225, show the greatest deviations in read depth between males and females, and likely represent a region where recombination was first suppressed between the emerging Z and W chromosomes (Stratum I). Our previous linkage mapping with GBS markers [48, 52] also identified sex-linked markers in scaffold 127 (Fig. S2), however this region shows far fewer differences in female:male read depth while having higher polymorphism in females relative to males. As a result, this likely represents a region where recombination has been suppressed very recently, or remains partially incomplete (Stratum II).

The SDR region spans a total of ∼3.4 Mb, or ∼3.1 Mb when excluding the putatively chimeric regions, and this estimate is somewhat smaller than that of our previous estimation of ∼5.3 Mb [49]. This difference is likely due to the fact that our previous estimate was based on a male assembly and included non-aligned regions on chromosome 15 of *P. trichocarpa*. In *Salix purpurea*, a close relative of *S. viminalis*, the SDR is also located on chromosome 15, however it is much larger (>10 Mb) [59]. It has been suggested that these sex chromosomes share a common origin [59], although it remains unclear whether the SDR in these two species is in the same syntenic region. In order to test whether the SDR regions overlapped between the two species, we aligned our *S. viminalis* genome assembly to the *S. purpurea* assembly. We found that all scaffolds inferred to be part of the *S. viminalis* SDR aligned to the SDR region in *S. purpurea* (Fig. S3), suggesting a shared origin, albeit with several potential rearrangements between them.

### Two evolutionary strata on the S. viminalis sex chromosomes

It is possible to quantify divergence between the sex chromosomes by comparing d_N_ (a measure of non-synonymous divergence) and d_S_ (a measure of synonymous divergence) between males and females in the sex-linked region. To accurately estimate this divergence, we constructed 10x Genomics Chromium *de novo* assemblies using one individual of each sex. Fully phased diploid genotypes were obtained for 65.8% and 61.6% of the genome in our female and the male samples respectively. Similar phasing efficiency was also achieved for chromosome 15 (Fig. S4). Our results show significantly greater d_N_ and d_S_ between Stratum I and the genomic average in our female sample, but not in our male sample (female d_S_ p=0.00072; female d_N_: p =0.000077; male d_S_ p =0.65; male d_N_ p =0.25, based on Mann-Whitney U-test relative to the genome, Fig. 2), indicating detectable divergence between the Z and W in this region. When Stratum II is also included, the SDR does not show significant divergence in the female (female d_S_ p =0.89; female d_N_ p =0.061; male d_S_ p =0.99; male d_N_ p =0.94, Mann-Whitney U-test relative to the genome) despite the presence of sex-linked markers in this region (Fig. S2), reinforcing the conclusion that either recombination was suppressed very recently in this region, or is not yet entirely complete. d_N_ and d_S_ were also marginally significantly higher between the pseudo-autosomal region (PAR) and the genome in females (d_S_ p =0.0019, d_N_ p =0.0133, Mann-Whitney U-test), but not in males (d_S_ p =0.93, d_N_ p=0.94, Mann-Whitney U-test).

**Figure 2.**
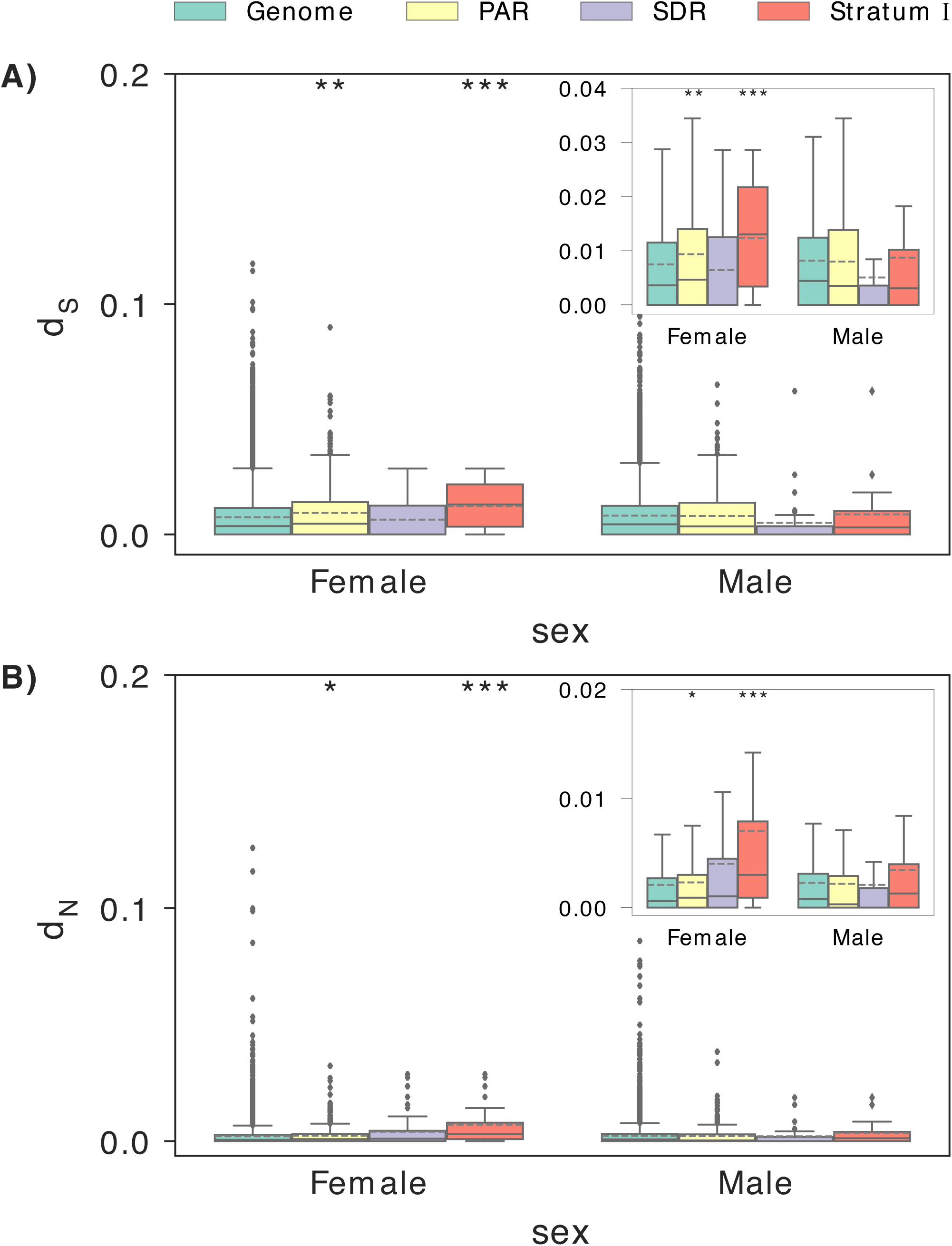
Comparison of polymorphisms at synonymous (d_S_) and non-synonymous (d_N_) sites. **A)** Boxplots of d_S_ estimates. **B)** Boxplots of d_N_ estimates. d_S_ and d_N_ were calculated based on the coding sequence alignment of phased diploid haplotypes from one female and one male individuals in the genome (excluding chromosome 15), the pseudo-autosomal region (PAR), the sex-determining region (SDR) and the more divergent Stratum I. The inset plots show the quartile distributions of d_S_ and d_N_ estimates without outliers. Significant values from Mann-Whitney U-test relative to the genome are indicated with asterisks: * p < 0.05; ** p < 0.01; *** p < 0.001.

Phylogenetic analysis of Z-W orthologs in conjunction with outgroup species can reveal the relative timing of recombination suppression [13]. We therefore used our phased male and female haplotypes in the SDR tohether with orthologous genes from two closely related *Salix* species (*S. suchowensis* and *S. purpurea*) and poplar (*P. trichocarpa*). Our phylogenetic analyses provide further support for two distinct evolutionary strata (Fig. 3, Fig. S5). Phylogenies based on genes located in Stratum I tend to show one female haplotype, corresponding to the W haplotype, clustering as an outgroup from the other three *S. viminalis* haplotypes (two male Z haplotypes and the female Z haplotype). This phylogenetic structure indicates that recombination ceased in Stratum I prior to *S. viminalis* speciation. The phylogenetic structure in Stratum II shows most female haplotypes clustered together with the male haplotypes, in line with more recent, or possibly partially incomplete, recombination suppression.

**Figure 3.**
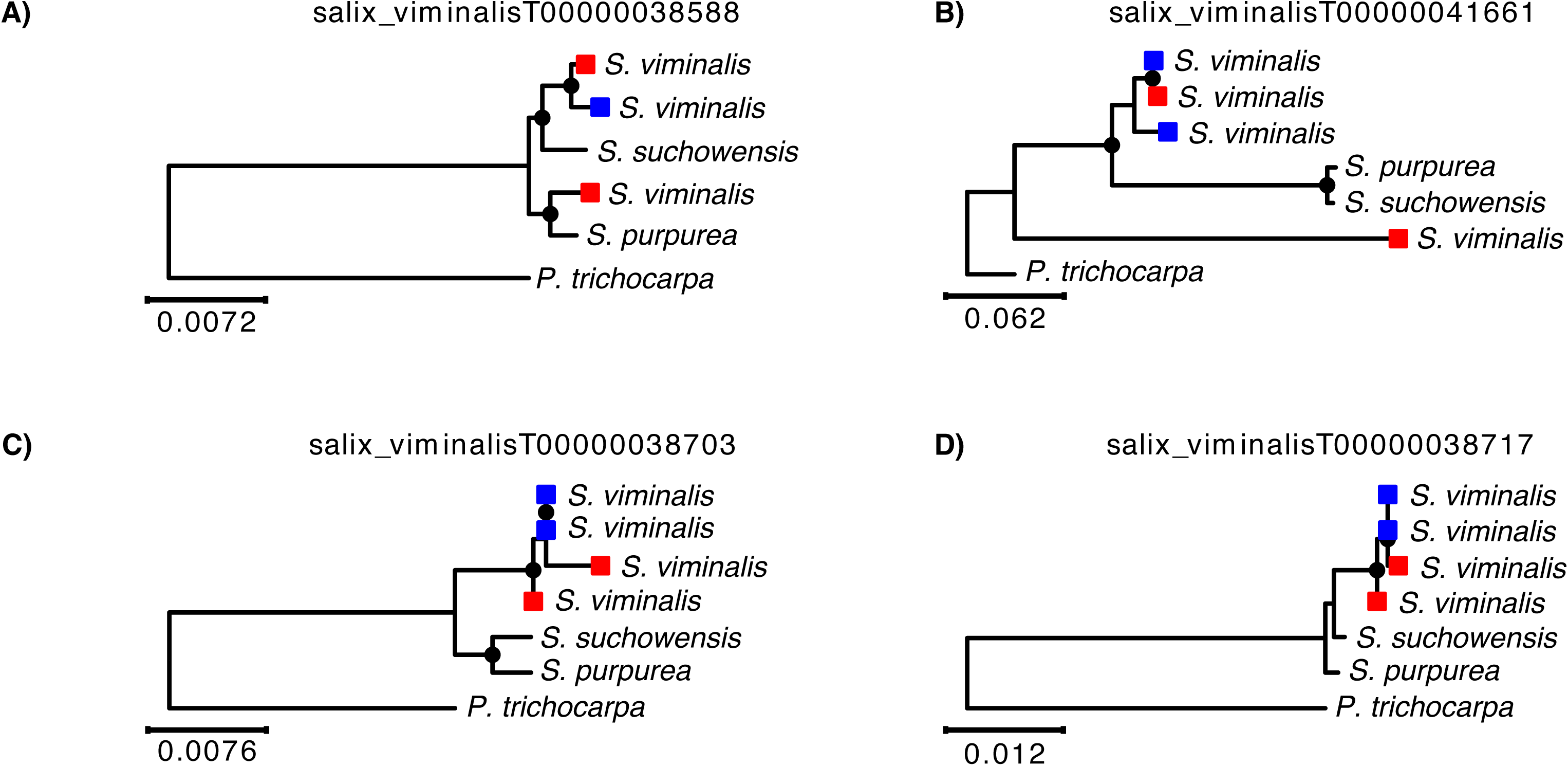
Examples of phylogenetic trees between gametologous gene pairs in the basket willow SDR. In panels **A)** and **B)**, the W-linked copy of the female gametolog is more divergent and does not cluster with the other *S. viminalis* haplotypes, indicating that suppression of recombination in Stratum I occurred prior to *S. viminalis* speciation. In panels **C)** and **D)** the female W-linked copy clusters within the species’ branch suggesting that recombination has been halted more recently. *S. viminalis* gametologs are indicated with red squares, male haplotypes are in blue. Trees were estimated by maximum likelihood. Bootstrap values >75% are indicated with black dots on the respective nodes. The poplar (*P. trichocarpa*) ortholog was used to root the trees.

Distinct evolutionary strata are evident in many sex chromosome systems [9–13], and the mechanism behind recombination suppression, whether it is a large-scale inversion on the sex-limited chromosome [18] or a more gradual suppression of recombination [14–17] remains unclear. Crucially, males and females can differ substantially both in frequency and in location of recombination hotspots [60–63], referred to as heterochiasmy. Local sex-specific recombination rates within the genome may be important in both initial sex chromosome divergence and subsequent expansion of the non-recombining region [26]. Importantly, once recombination has been halted around the SDR in the heterogametic sex, selection to maintain gene order is abolished [64], and selection against inversions is greatly reduced. This suggests that inversions might follow recombination suppression, as has been recently observed [65], even if they are not the cause of recombination suppression initially.

It is worth noting that we observed considerable overlap in both d_S_ and d_N_ estimates (Fig.2) between the two strata and also the incomplete segregation of some female Stratum I Z and W haplotypes (Fig. S5), suggesting a gradual divergence in the sex chromosomes of *S. viminalis*. This gradual divergence is not consistent with a major inversion, which would result in a more similar phylogenetic signal for all Z-W orthologs within the inversion as recombination would be suppressed at the same time. Older sex chromosomes also show substantial variation in divergence within perceived strata [10, 13], however the limited number of loci remaining on the oldest regions of sex-limited chromosome complicates these analyses. In these older systems, strata may also have formed through shifts in sex-specific recombination hotspots, resulting in gradual expansions rather than large-scale inversion events.

Furthermore, if inversions are the cause of recombination suppression between the Z and W, we would expect our female assembly to be heterozygous for inversions between the Z and W chromosomes in strata. We note that we observe no evidence of large-scale inversions associated with either Stratum I or Stratum II in our assembly. It is of course possible that inversions formed within the few remaining breakpoints in between our scaffolds, which we would not be able to detect. However, the long-read nature of our assembly, and the resulting large contig size, offer substantial power to detect such inversions, reducing the likelihood of type II error.

Together, our evidence suggests that at the earliest stages of sex chromosome formation and expansion, selection against recombination is a gradual process, and may result from changes in sex-specific recombination hotspots. Therefore, theoretical models about local changes in heterochiasmy as a result of sexually antagonistic alleles [62, 63] may prove to be key to sex chromosome evolution. Alternatively, recent evidence from fungal mating-type chromosomes, analogous to sex chromosomes in many ways, has suggested non-adaptive explanations for the origin and expansion of the non-recombining region [66, 67].

This model also explains some of the curious intra-specific heterogeneity in the extent of sex chromosome divergence in younger systems [32–35]. If recombination suppression occurs more gradually, population-level differences in sex-specific recombination hotspots, often observed [61], will drive different levels of divergence in the earliest stages of sex chromosomes.

### Degeneration of the W chromosome

Although studies of old, highly degenerate Y and W chromosomes have revealed the accumulation of significant repetitive DNA [68, 69], it remains unclear how quickly this material accumulates after recombination suppression. Additionally, the build-up of repetitive elements on the W chromosome may in itself act as a mechanism to suppress recombination with the corresponding region of the Z [70–72]. However, the difficulty associated with phasing short read data has previously hampered efforts to study the earliest stages of sex chromosome divergence. Although it is possible to identify sex-specific transcripts from pedigrees based on inheritance through familial pedigrees [73–76], this method misses non-coding sequence, making it difficult to assess whether non-coding repetitive elements are associated with the earliest stages of recombination suppression.

In order to identify W-specific sequence, we mapped female and male re-sequencing reads to our female assembly. We were able to identify an additional subset of 35 scaffolds spanning ∼3.3 Mb and with 119 protein coding genes (Table S4), that likely represent W-specific sequence, i.e., with significant excess of female:male read coverage over the entire scaffold length based on genomic confidence intervals. Despite the recent origin of recombination suppression, these scaffolds show a significant enrichment of repetitive sequences in comparison with both the corresponding Z-linked portion of the SDR and the genomic average (Fig. S6, W-genome p < 1×10^-46^; W-SDR p = 0.00058, Mann-Whitney U-test). These results suggest that either repetitive sequence can accumulate very quickly following the arrest of recombination, or alternatively repetitive elements may in fact act to halt recombination in the absence of inversions.

The loss of recombination on the sex-limited SDR has important evolutionary effects, namely the build-up of deleterious variation and repetitive elements, as well as loss of gene activity [6–8]. The latter effect in particular can lead to profound differences in gene content between X and Y or Z and W chromosomes in older sex chromosome systems [6]. Studies in other plant sex chromosomes have indicated that gene loss occurs in the SDR [8, 47], however it remains unclear how quickly this occurs. Additionally, the extended haploid phase in plants may prevent loss of SDR genes expressed in the haploid phase [15, 46].

In order to identify gene content differences between the Z and the W chromosome, we used two of the W-linked scaffolds identified above, scaffolds 148 and 211. These scaffolds align almost entirely to the SDR where read mapping coverage is male-biased (Z-linked), as would be expected for sex-linked homologous regions (Fig. 4A). In both cases we observed a high degree of synteny in the aligned regions, indicating that both gene content and gene order are still largely conserved between Z and W homologs, even in the most divergent region of the SDR (Fig. 4B, 4C). This is likely a function of both the recent divergence of this sex chromosome system [49], as well as the preservative effects of haploid selection on genes expressed in plant reproductive tissues. Nevertheless, seven protein coding genes on the corresponding Z-linked scaffolds with known products are missing from the W assembly. Using a translated BLAST search of these proteins to the corresponding Z-linked scaffolds and considering a minimum query coverage of 80%, we inferred that at least two of them (os02g0180000 on scaffold 163 and TIR on scaffold 225) have likely been pseudogeneized on the W. These results suggest that gene loss can occur very quickly, even in nascent sex chromosome systems.

**Figure 4.**
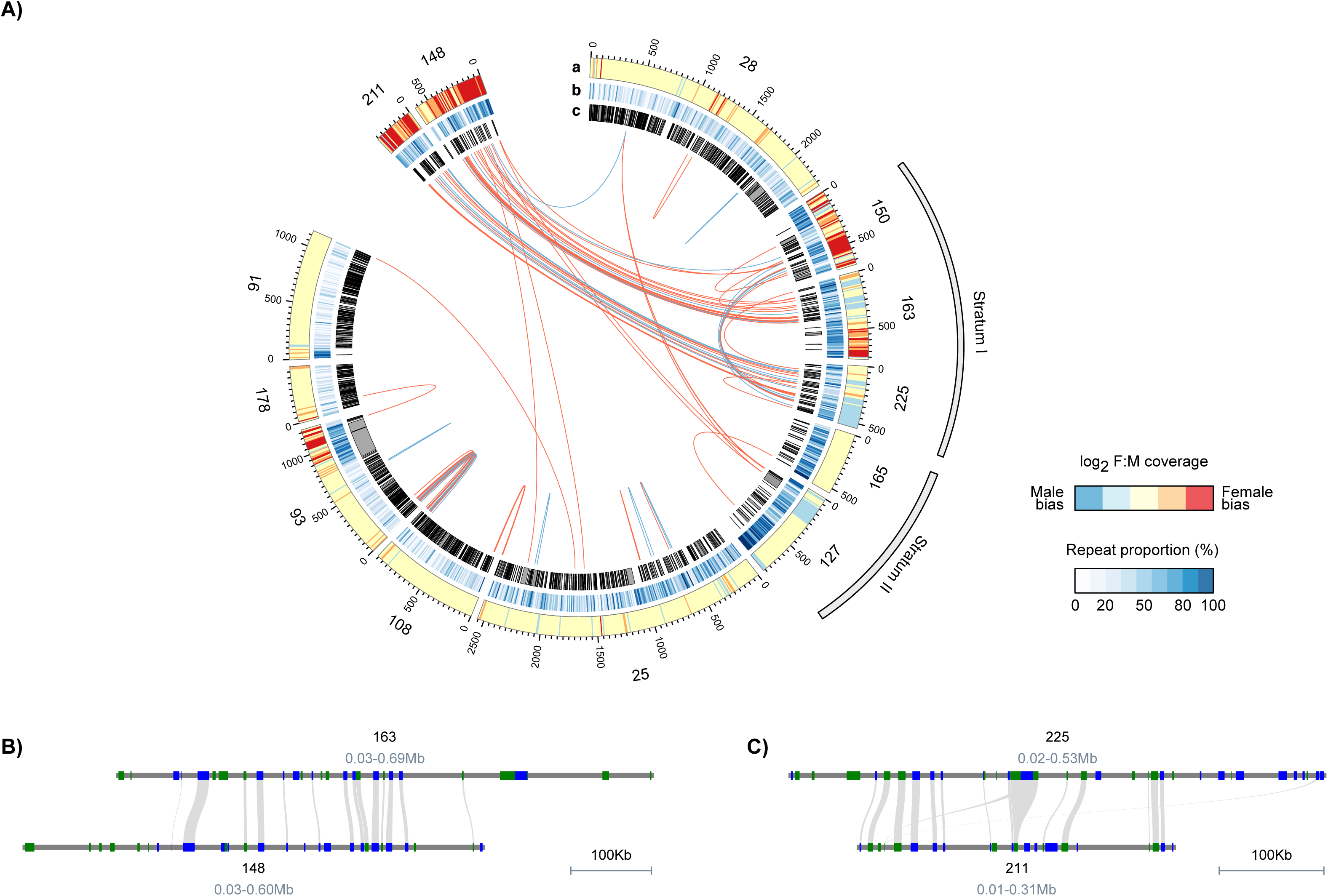
Synteny analyses of Z- and W-linked resolved haplotypes. **A)** Circular plots showing that scaffolds 148 and 211 are W-linked and align to the SDR of chromosome 15. From the outside to the center, (a) depicts the heatmap of log_2_ females: males read depth in non-overlapping windows of 5 kb, (b) shows the repeat proportion in non-overlapping windows of 10 kb and (c) indicates the location of annotated genes. Links between genes were computed from the best BLASTP hits and are colour coded relative to the BLASTP alignment percent identity, with percent identity >80% in blue and >90% in red. Positions are shown in kb. **B)** and **C)** reveal a highly conserved synteny between Z- and W-linked scaffolds.

We also scanned for genes unique to the *S. viminalis* W chromosome, or without preserved synteny to the Z homolog, as possible sex determining loci. We recovered several genes, including WOX1, as well as two genes in tandem of the two-component response regulator, ARR5 and ARR17 (Table 1). In *Arapidopsis thaliana*, ARR proteins are the final targets of the cytokinin signalling system, which is known to play important roles in flower development and floral sex differentiation [77, 78]. WOX1 is a WUSCHEL-related homeobox protein, involved in the central regulatory pathway that coordinates stem cell proliferation with differentiation [79]. Interestingly, the *Silene latifolia* homolog of WOX1, SlWUS1, is also sex-linked on the X chromosome and apparently lost the homologous copy in the Y chromosome [80]. We found a WOX1 ortholog in scaffold 150 with ∼88% sequence identity and an ortholog for ARR5 on scaffold 28 with ∼95% sequence identity suggesting either recent duplications or translocations to the W-linked sequence. For ARR17, we did not recover an ortholog in the genome, or evidence for a pseudogene in the Z chromosome, suggesting that it most likely originated through a translocation to the W.

**Table 1.** Genes on W chromosome scaffolds 148 and 211 with non-preserved synteny relative to the homologous region on the Z chromosome. Orthologs were searched with BLASTP using an e-value threshold of 1 × 10^-3^ and 75% minimum sequence identity.

| Scaffold | Gene | Product | Scaffold of best ortholog (location in *P. trichocarpa*) |
| --- | --- | --- | --- |
| 211 | ADT2 | Arogenate dehydratase/prephenate dehydratase 2, chloroplastic | 100 (Chr08) |
| 211 | 30220 | hypothetical protein |  |
| 211 | 30217 | hypothetical protein |  |
| 211 | POPTR_0012s05040g | L-Ala-D/L-amino acid epimerase |  |
| 211 | 30210 | hypothetical protein | 71 (Chr18) |
| 211 | FBA | Fructose-bisphosphate aldolase | 402* (Chr15) |
| 148 | KP1_5 | Kinesin KP1 | 150 (Chr15) |
| 148 | ESP3_4 | Pre-mRNA-splicing factor ATP-dependent RNA helicase DEAH1 | 127 (Chr15) |
| 148 | CDC48MEE29 | Cell division cycle protein 48 homolog | 47 (Chr12) |
| 148 | ESP3_2 | Pre-mRNA-splicing factor ATP-dependent RNA helicase DEAH1 | 28 (Chr15) |
| 148 | ESP3_6 | Pre-mRNA-splicing factor ATP-dependent RNA helicase DEAH1 | 127 (Chr15) |
| 148 | ARR5_2 | Two-component response regulator ARR5 | 25 (Chr15) |
| 148 | ARR17 | Two-component response regulator ARR17 |  |
| 148 | WOX1_4 | WUSCHEL-related homeobox 1 | 150 (Chr15) |
| 148 | ATM_6 | Serine/threonine-protein kinase ATM | 25 (Chr15) |
| 148 | BADH4_2 | Betaine aldehyde dehydrogenase, chloroplastic | 326 (Chr12) |
| 148 | ZDS_7 | Zeta-carotene desaturase, chloroplastic/chromoplastic | 593 (Chr15) |
| 148 | 27648 | hypothetical protein |  |
| 148 | CDKE-1_12 | Cyclin-dependent kinase E-1 | 4 (Chr01) |
| 148 | 27660 | hypothetical protein |  |

\* scaffold 402 was inferred as an allelic variation of scaffold 150.

It is worth noting that dioecy evolved early in the Salicaceae lineage in which *S. viminalis* is embedded, and is shared by most members of the clade [81]. This means that the standard model for the evolution of sex chromosomes in plants, which assumes an immediate hermaphrodite ancestor, may not be applicable. The model posits two linked mutations encoding male-and female-sterility [82] as the progenitor of sex chromosomes, and this model has received some empirical support [41]. However, the ancient dioecy found in Salicaceae and the observation of small and heterogeneous levels of divergence in the basket willow [49] and poplar [37] sex chromosomes are difficult to reconcile with this two-gene model. Indeed, recent work has pointed out alternative sex determination mechanisms in flowering plants, either determined by a single gene as in the case of *Diospyros* [40] or, as in *Cucumis*, as a polygenic trait controlled by several genes distributed across different chromosomes [83]. The Salicaceae family with its young sex chromosomes derived from ancient dioecy therefore provides a valuable comparative system to elucidate this process.

### Concluding remarks

Here, we use multiple types of single-molecule sequencing to assemble the genome of the basket willow *S. viminalis*, and used this to reveal the earliest stages of sex chromosome evolution. This approach allows us unprecedented power to phase our data, allowing us to resolve Z and W haplotypes at this early stage of divergence. Our results suggest that the SDR is of limited size and divergence, and we recover no evidence that recombination suppression is due to a large-scale inversion. Even at this early stage of divergence, we see evidence of pseudogenization and the accumulation of repetitive elements in the SDR, suggesting that these processes occur very swiftly after recombination ceases. In total, our results shed new light on the fundamental process of sex chromosome formation.

## Materials and Methods

### Plant material and DNA extraction

Fresh young leaves (approximately 200 mg) were sampled from a female and a male *S. viminalis* (accession 78183 and 81084, respectively), described in [51] and [84] and DNA was extracted following a CTAB-protocol described in [49]. In brief, approximately 200 mg fresh leaves were snap frozen and pulverized. To every sample, 950 μl of extraction buffer (100 mM TrisHCl pH 7.5–8, 25 mM EDTA, 2 M NaCl, 2% (w/v) CTAB, 2% (w/v) PVP K30, 5% (w/v) PVPP, 50 μg/ml RNAse) was added and the sample was thoroughly mixed before incubation for 30 min at 65 °C. Subsequently, 300 μl Chloroform:isoamylalcohol 24:1 was added, the sample mixed and centrifuged for 10 min at 13,000 rpm, the supernatant was transferred to a new tube, and the process repeated. 1.5 volumes of ice-cold isopropanol was added to the supernatant followed by an incubation over night at −20 °C. After centrifugation for 10 min at 13,000 rpm at 4 °C, the supernatant was removed and the pellet rinsed with chilled 100% EtOH followed by another centrifugation of 5 min at 13,000 rpm at 4 °C. The supernatant was then removed and the DNA was air dried before it was dissolved in 100 μl TE buffer (10 mM TrisHCl, 1 mM EDTA). DNA concentration was assessed by Qubit 3.0 Fluorometer (Thermo Fisher Scientific).

### PacBio long-read library preparation and sequencing

A single SMRT-bell library with 20 kb insert size was constructed from 10 µg of pure high-molecular weight DNA from one *S. viminalis* female (accession 78183) according to the manufacturer’s protocol (Pacific Biosciences). This library was sequenced on 48 SMRT cells using P5-C3 chemistry and 4 hour movies were captured for each SMRT cell using the PacBio RSII sequencing platform (Pacific Biosciences). Primary analysis and error correction of the raw data was done using SMRT Portal (Pacific Biosciences). After filtering, the mean read length was 8,924 bp (longest read was 61 kbp) and a total of ∼19.2 Gbp of data were recovered.

### 10x Genomics Chromium linked reads library preparation and sequencing

For both accessions (78183 and 81084) sequencing libraries were prepared from 0.75 ng DNA using the Chromium TM Genome Library preparation kit according to the CG00022_Chromium Genome Reagent Kit User Guide_RevA. The library preparation was performed according to the manufacturers’ instructions with the exception that 0.75 ng was used for library preparation instead of 1.25 ng recommended by the manufacturer’s instructions. This was done to account for the smaller genome size of *S. viminalis* compared to the human genome for which the protocol was optimized. The libraries were sequenced on an Illumina HiSeqX with a paired-end 150bp read length using v2.5 sequencing chemistry (Illumina Inc.), resulting in ∼58 Gb of data

### DNA extraction and short-read Illumina sequencing

We generated additional Illumina sequencing data for the female accession 78183, the same accession used to assemble the reference genome. DNA was extracted from fresh leaves using the Fast DNA Kit (MP Biomedicals) according to the manufacturer’s instructions. Two libraries with 165 and 400 bp insert size respectively were generated with the TruSeq DNA v2 kit (manual #15005180) following the manufacturers protocol and sequenced on one lane each with Illumina HiSeq2000, 100bp paired-end read length and v3 chemistry generating ∼28 GB of bases (Table S1).

### Reference genome assembly and annotation

Falcon v0.4.2 [85] was used to assemble the sub-reads from 48 SMRT cells. This first draft assembly was then polished using Quiver from the Pacific Biosciences’ SMRT suite (v2.3.0) with the PacBio reads. The resulting assembly was then corrected with Pilon v 1.17 [86] using both Illumina libraries from the same individual at 80X and 53X coverage. In addition, a 10x Genomics assembly for the same female individual was also obtained using the pseudohap-style output of Supernova v2.0.1 [87]. This 10x Genomics assembly and the PacBio assembly were then merged using Quickmerge v20160905 [88], increasing the assembly size by ∼8 Mb. Finally, the preads (corrected PacBio reads obtained after the first step of Falcon assembly) and the Supernova pseudohap assembly were used to scaffold the merged assembly using LINKS v1.8.4 [89]. Finally, we corrected some homozygous SNPs and small insertions and deletions in the assembly using Long Ranger v2.1.2 with the 10x Genomics Chromium reads of the same female individual.

Annotation of the *S. viminalis* reference genome was performed with Maker v3.00.0 [54]. The Maker pipeline was run twice; first based on protein and RNA sequence data only (later used to train *ab-initio* software) and a second time combining evidence data and *ab-initio* predictions. High-confidence protein sequences were collected from the Uniprot database [90], for proteins belonging to the Swissprot section that contain only manually annotated and reviewed curations (downloaded on August 2016), and two other specific protein sets from *Salix suchowensis* and *Populus trichocarpa*. Furthermore, to support gene predictions we also used selected libraries of RNA-seq data from our previous studies collected from different individuals and tissues [49, 55]. As basis for the construction of gene models, we combined *ab-initio* predictions from three sources (Augustus v2.7 [91], GeneMark_ES_ET v4.3 [92] and SNAP [93]). GeneMark_ES_ET was self-trained with the genome sequence. To train Augustus and SNAP, we first ran the Maker pipeline a first time to create a profile using the protein evidence along with RNA-seq data. Both Augustus and SNAP were then trained with a selected set of genes from this initial evidence-based annotation. We excluded genes with an Annotation Edit Distance (AED) score equal to 1 to avoid potentially false annotations. Functional inference for genes and transcripts was performed using the translated CDS features of each coding transcript. Protein sequences were searched with BLAST in the Uniprot/Swissprot reference dataset in order to retrieve gene names and protein functions as well as in the InterProscan v5.7-48 database to retrieve additional annotations from different sources.

We created a repeat library with an in-house pipeline using RepeatModeler v1.0.8 [94]. Identification of repeat sequences in the genome was performed using RepeatMasker v4.0.3 [95] and RepeatRunner [96]. tRNAs were predicted with tRNAscan v1.3.1 [97] and broadly conserved ncRNAs were predicted with the Infernal package [98] using the RNA family database Rfam v11 [99].

### Identification of allelic scaffolds in single-molecule de-novo assemblies

Linked reads for the female and male accessions were assembled with Supernova v2.0.1 [87]. Fully phased heterozygous haplotypes, together with non-phased sequence (nominally homozygous), were obtained using the megabubbles-style output and a minimum sequence length of 1 kb. Diploid assemblies were soft-masked with RepeatMasker v4.0.7 [95] with the “RMBlast” v2.6.0+ search engine and using our custom *S. viminalis* repeat library generated during genome annotation.

We used sequence alignments in order to identify homologous haplotypes in our single-molecule assemblies. A repeat-masked assembly is first aligned to itself with LAST v926 [100] using the sensitive DNA seeding MAM4 [101] and masking of repeats during alignment with the -cR11 option. To avoid false matches caused by repetitive sequences and paralogous scaffolds, orthologous alignments were generated with last-split and alignments mostly comprised of masked sequence were then discarded with last-postmask. Scaffolds were considered to represent allelic variants in the assembly if the overlap exceeded 25% of sequence length after repeat masking, and with sequence identity > 80% to other longer scaffolds.

### *Anchoring scaffolds to* Populus trichocarpa

Pairwise alignments between *P. trichocarpa* v10.1 (downloaded from PopGenie v3 [56]) and our *S. viminalis* assembly were generated from repeat-masked genomic sequence using LAST v926 [100]. We first prepared an index of the poplar genome using the sensitive DNA seeding MAM4 [101], using the masking repeat option -cR11 during alignment. A suitable substitution and gap frequencies matrix was then determined with last-train, using parameters --revsym --matsym --gapsym -C2. Alignments were made with lastal, using the parameters -m100 -C2 followed by last-split –m1 to find 1-to-many willow-poplar orthologous matches. Finally, alignments that were composed primarily of masked sequence were discarded with last-postmask. One-to-one willow-poplar alignments were made by swapping both sequences and repeating the orthology search as above.

Neighboring alignments with <10 kb gap lengths were linked into a single path and the longest tiling path was used to assign scaffolds to poplar chromosomes. Forward or reverse scaffold orientation relative to poplar chromosomes was similarly obtained requiring that the total length of one alignment direction was >70% compared to the other orientation, otherwise the original orientation was kept. If the longest tiling path for a particular scaffold did not agree with its overall alignment path on the poplar chromosome, the scaffold was marked as unlocalized.

### Preprocessing of Illumina reads

Whole-genome DNA sequencing reads were quality assessed with FastQC v0.11.5 [102] and preprocessed with BBTools v37.02 “bbduk” [103] to remove adapter sequences, trim regions with average quality scores below Q10 from both ends of reads and to filter out reads aligning to PhiX-174 genome (a commonly used spike-in control in Illumina sequencing runs). After filtering, read-pairs were excluded from downstream analyses if either read had an average quality score <Q20 or was <50 bases in length. The same criteria of quality assessment and filtering were used for RNA-seq data.

### Coverage and polymorphism analysis

Alignments to the genome assembly were performed with BWA v0.7.15-r1140 using the MEM [104] algorithm and default options. General processing of SAM/BAM files was performed with SAMtools v1.6 [105] and duplicated reads were flagged with biobambam v2.0.72 [106] after alignment. Per-site coverage was computed with the SAMtools depth command after filtering out reads with mapping quality ≥ Q3 that map to multiple locations, reads with secondary alignments and duplicated reads. We then calculated the effective coverage value per scaffold and in non-overlapping windows of 10 kb, as the mean per site coverage of every site in that class. To account for differences in the overall coverage between individuals, the coverage data were normalized for the median coverage value of each individual in the respective class.

Polymorphism analyses were conducted using the same filters as above. Read alignments were then converted to nucleotide profiles with the sam2pro program of mlRho [107]. Only sites with a per-site coverage ≥5 and a SNP called for bi-allelic sites with a minor allele frequency ≥30% within an individual were analysed. The average SNP density per scaffold, and window, was calculated as the number of SNPs divided by the number of sites that passed the coverage threshold of ≥5 for the respective class.

In order to avoid infinitely high numbers associated with log_2_ 0 when calculating the log_2_ difference of coverage or SNP density between females and males, we added a small number (0.1) to each value. The 95% confidence intervals for the sliding window distributions were estimated from the mean bootstrap values with resampling of 1,000 random sets of 25 windows from autosomes. We excluded the entirety of chromosome 15 (the sex chromosome), including the PAR, in the bootstrapping procedure to avoid potential linkage effects resulting from the SDR.

To identify potentially W-linked scaffolds in the assembly, we proceeded as above and calculated the log_2_ F:M coverage differences for each scaffold. All scaffolds where the normalized female coverage was <10% of the normalized whole-genome coverage were excluded. This is a conservative approach because of the difficulty associated with mapping to highly repetitive potential W-linked scaffolds. These scaffolds are therefore likely to remain undetected. Scaffolds were considered W-linked if the log_2_ F:M coverage difference was >95% the genome average.

### Quantification of gene expression

Preprocessed RNA-seq reads [49, 55] were filtered for rRNA using Bowtie v2.3.2 [108] and the SILVA release 128 database of LSU and SSU NR99 rRNAs [109]. Filtered reads were then aligned to the reference assembly using HISAT2 v2.1.0 [110] with options --no-mixed --no-discordant. The resulting alignments for each library were sorted and merged by individual and by tissue (catkin and leaves) with SAMtools v1.6 [105]. Read counting per gene was performed using the count command of HTSeq [111] and reads per kilobase mapped (rpkm) expression values were calculated with edgeR [112]. Only genes with an rpkm ≥1 in at least one sample were considered in further analyses.

### Annotation lift-over to 10x Genomics diploid assemblies

Our reference genome annotation was transferred independently for each of the inferred haplotypes derived from our 10x Genomics de-novo assemblies of female and male genomes using UCSC Genome Browser’s utilities [113]. First, a pairwise alignment between each haplotype and the non-redundant reference genome was generated as described above with LAST v926 [100]. Alignments were then converted into a series of syntenic chains and nets, tuned for more divergent genomes (axtChain -linearGap=loose), using the same scoring matrix generated during the LAST alignments. Finally, annotations were moved to the haplotype assemblies using the liftOver utility with a minimum 75% ratio of mapped bases between features. Only the longest isoform of each gene was considered in the lift-over. With this approach, we transferred ∼25,159 genes per diploid haplotype or ∼80% of the complete annotation.

We further attempted to recover additional genes not lifted initially by aligning each gene individually back to the haplotype assemblies with BLAT v170523 [114], (-minIdentity=30 - minScore=12 -stepSize=5 -repMatch=2253 -extendThroughN), keeping the highest scoring alignment for each query. In order to avoid potential problems caused by the BLAT alignment of paralogous sequences, we counted the average number of haplotypes aligned to each reference gene (for a fully phased diploid region we expect 2 haplotypes). These counts were then bootstrapped with 1,000 iterations all alignments for which the haplotype coverage was below the lower bootstrap 95% confidence interval (∼1.6X coverage) were excluded. This procedure recovered an average of 364 additional genes per haplotype.

### Divergence analysis of diploid genotypes

We calculated rates of divergence at synonymous (d_S_) and non-synonymous (d_N_) sites between the coding sequences of diploid genotypes for each sex separately. Only sequences with a valid start codon, without internal stop codons and with a minimum sequence length of 120 bases were analysed. After this initial filter, pairwise alignments for the two haplotypes were obtained with PRANK v140603 [115] and d_S_ and d_N_ estimates were calculated using the method of Yang and Nielsen [116] as implemented in the yn00 program of PAML v4.9h [117]. Pairwise comparisons with d_S_ > 0.2 were excluded, thereby avoiding the incorrect assignment of orthologs.

### Phylogenetic analysis

We use gene trees to determine the relative age of recombination suppression for the haplotypes in each identified sex chromosome strata. In addition to our non-redundant *S. viminalis* genome, coding sequences for *S. suchowensis* v4.1 and *P. trichocarpa* v10.1 were obtained from PopGenie v3 [56] and sequences for *S. purpurea* v1.0 were obtained from Phytozome v12 [118]. Only the longest transcripts were considered. We first use the Conditional Reciprocal Best BLAST method [119], with a BLAST e-value cut-off < 1×10^-5^, to identify 14,255 one to one orthologs across all four species (*S. viminalis, S. suchawensis, S. purpurea and P. trichocarpa*). For each ortholog group, we searched for the *S. viminalis* homolog in the lifted annotation of the female and male phased diploid assemblies and aligned all species’ sequences with MAFFT v7.313 [120]. Aligned columns with > 40% gaps and taxa with > 40% of missing data were removed. Maximum likelihood phylogenetic trees were obtained with RAxML v8.2.12 [121] using the rapid bootstrap algorithm with 100 bootstraps and the GTRGAMMA model of sequence evolution. Trees were rooted on the *P. trichocarpa* branch and were only considered if the two female haplotypes were present. Phylogenetic tree analyses were performed with ETE3 [122].

## Data accessibility

Genome sequencing data and annotation generated for this study have all been deposited in EBI’s ENA (https://www.ebi.ac.uk/ena) under project number PRJEB31619. IPython notebooks and additional data necessary to reproduce the main figures are available from Dryad Digital Repository doi:XXXXXX.

## Acknowledgments

This work was funded by the European Research Council (grant agreements 260233 and 680951) to JEM. JEM also gratefully acknowledges further support from a Royal Society Wolfson Merit Award and a Canada 150 Research Chair. We acknowledge the use of the University College London Legion High Performance Computing Facility (Legion@UCL), and associated support services, in the completion of this work. Sequencing and annotation were funded by the Swedish Energy Agency, grant 30599-5 and from the Swedish Research Council for Environment, Agricultural Sciences and Spatial Planning (Formas) grant 2016-20031. We also acknowledge the support provided by NBIS (National Bioinformatics Infrastructure Sweden). Sequencing was performed by the National Genomics Infrastructure (NGI) Sweden at Science for Life Laboratory. NGI is supported by the Swedish Research Council and the Knut and Alice Wallenberg Foundation, which supported AC, BN, and EP-W during the course of this work. We thank A. Corral-Lopez, I Darolti, B. Furman, D. Metgzer, B. Sandkam, J. Shu and W. van der Bijl for helpful comments and suggestions.

