## Supplementary material for "Single-molecule genome assembly of the Basket *Willow, Salix viminalis*, reveals earliest stages of sex chromosome expansion"

**Table S1.** Whole-genome DNA sequencing data used in this study.

| Accession | Sex | Technology | Number of reads (millions) | Number of Megabases (Gb) | Reference |
|---|---|---|---|---|---|
| 78183 | Female | Pacific Biosciences | 2.7 | 19 | This study |
| | | 10x Genomics | 390 | 59 | This study |
| | | Illumina – 165 bp insert | 191 | 19 | This study |
| | | Illumina – 400 bp insert | 289 | 28 | This study |
| 78021 | Female | Illumina | 204 | 26 | Pucholt et al. 2017 |
| 78195 | Female | Illumina | 199 | 20 | Pucholt et al. 2017 |
| 81084 | Male | 10x Genomics | 429 | 64 | This study |
| | | Illumina | 210 | 21 | Pucholt et al. 2017 |
| T76 | Male | Illumina | 214 | 27 | Pucholt et al. 2017 |

**Table S2.** Assembly statistics for the full and non-redundant assemblies of *Salix viminalis*. DNA- and RNA-seq mapping were performed with read data from the same individual used in the assembly.

| Metric | Full assembly | | Non-redundant assembly | |
|---|---|---|---|---|
| Number of scaffolds | 2,372 | | 1,467 | |
| Total bases (bp) | 357,061,245 | | 303,195,481 | |
| Longest sequence (bp) | 7,291,771 | | 7,291,771 | |
| N50 | 73 | | 54 | |
| L50 (bp) | 1,314,944 | | 1,658,965 | |
| N90 | 434 | | 203 | |
| L90 (bp) | 77,063 | | 300,484 | |
| % gaps (N) | 0.73 | | 0.48 | |
| BUSCO - genome | 79.5% single copy | | 82.3% single copy | |
| | 15.0% duplicated | | 10.0% duplicated | |
| BUSCO - proteins | 73.1% single copy | | 75.2% single copy | |
| | 14.1% duplicated | | 9.7% duplicated | |
| DNA-seq mapping | 98.41% (mapped) | 90.45% (properly paired) | 97.72% (mapped) | 89.31% (properly paired) |
| RNA-seq mapping (2 tissues) | 84.28% (mapped) | 71.14% (properly paired) | 81.85% (mapped) | 69.16% (properly paired) |

**Table S3.** Characterization of the different annotation classes for the *Salix viminalis* assembly.

| Source | Value |
|---|---|
| Gene number | 36,490 |
| Gene span | 173,927,686 |
| Gene average length (stdev) | 4,766 (10,540) |
| Genes with functional annotation (%) | 28,212 (77.3%) |
| Transcript number | 49,131 |
| Transcript average length (stdev) | 6,485 (16,044) |
| Intron number | 294,271 |
| Intron average length (stdev) | 786 (4,636) |
| 3’ UTR number | 26,926 |
| 3’ UTR average length (stdev) | 404 (423) |
| 5’ UTR number | 26,883 |
| 5’ UTR average length (stdev) | 245 (307) |
| Exon number | 343,402 |
| Exon average length (stdev) | 254 (341) |
| CDS number | 49,131 |
| CDS average length (stdev) | 298 (403) |
| Genes with multiple transcripts (%) | 6,158 (16.9%) |
| Average number of transcripts per gene | 1.35 |
| Average number of exons per transcript | 6.99 |
| Single exon transcripts (%) | 4,367 (8.9%) |
| Gene coverage | 48.7% |
| CDS coverage | 4.1% |
| ncRNA | 3,469 |
| tRNA | 1,139 |

**Table S4.** List of all genes found on putatively W-linked scaffolds.

| Scaffold | Gene | Product |
|---|---|---|
| 211 | CCT4_2 | T-complex protein 1 subunit delta |
| 211 | GRDP1 | Glycine-rich domain-containing protein 1 |
| 211 | ADT2 | Arogenate dehydratase/prephenate dehydratase 2, chloroplastic |
| 211 | 30233 | hypothetical protein |
| 211 | AT1g18030_2 | Probable protein phosphatase 2C 8 |
| 211 | AT1g22950_3 | Uncharacterized PKHD-type hydroxylase At1g22950 |
| 211 | PAPS1_2 | Nuclear poly(A) polymerase 1 |
| 211 | 30222 | hypothetical protein |
| 211 | GGR_2 | Heterodimeric geranylgeranyl pyrophosphate synthase small subunit, chloroplastic |
| 211 | 30220 | hypothetical protein |
| 211 | 30219 | hypothetical protein |
| 211 | HIP1 | Probable E3 ubiquitin-protein ligase HIP1 |
| 211 | 30217 | hypothetical protein |
| 211 | 30215 | hypothetical protein |
| 211 | RTNLB9 | Reticulon-like protein B9 |
| 211 | POPTR_0012s05040g | L-Ala-D/L-amino acid epimerase |
| 211 | NACK1_2 | Kinesin-like protein NACK1 |
| 211 | 30210 | hypothetical protein |
| 211 | 30206 | hypothetical protein |
| 211 | AMSH1_2 | AMSH-like ubiquitin thioesterase 1 |
| 211 | 30204 | hypothetical protein |
| 211 | FBA | Fructose-bisphosphate aldolase |
| 148 | KP1_5 | Kinesin KP1 |
| 148 | ESP3_4 | Pre-mRNA-splicing factor ATP-dependent RNA helicase DEAH1 |
| 148 | CDC48MEE29 | Cell division cycle protein 48 homolog |
| 148 | ESP3_2 | Pre-mRNA-splicing factor ATP-dependent RNA helicase DEAH1 |
| 148 | ESP3_6 | Pre-mRNA-splicing factor ATP-dependent RNA helicase DEAH1 |
| 148 | ARR5_2 | Two-component response regulator ARR5 |
| 148 | ARR17 | Two-component response regulator ARR17 |
| 148 | WOX1_4 | WUSCHEL-related homeobox 1 |
| 148 | CML27_4 | Probable calcium-binding protein CML27 |
| 148 | GC2 | Golgin candidate 2 |
| 148 | ATM_6 | Serine/threonine-protein kinase ATM |
| 148 | BADH4_2 | Betaine aldehyde dehydrogenase, chloroplastic |
| 148 | ZDS_7 | Zeta-carotene desaturase, chloroplastic/chromoplastic |
| 148 | HMGIY2_2 | HMG-Y-related protein A |
| 148 | MPK15 | Mitogen-activated protein kinase 15 |
| 148 | AT4g28100_3 | Uncharacterized GPI-anchored protein At4g28100 |
| 148 | 27648 | hypothetical protein |
| 148 | MYB6 | Transcription repressor MYB6 |
| 148 | CDKE-1_12 | Cyclin-dependent kinase E-1 |

| | | |
|---|---|---|
| 148 | AIP1-2 | Actin-interacting protein 1-2 |
| 148 | BGLU44_3 | Beta-glucosidase 44 |
| 148 | EXO1_2 | Exonuclease 1 |
| 148 | MYBL | Myb-like protein L |
| 148 | GB1_3 | Guanine nucleotide-binding protein subunit beta-like protein |
| 148 | PEX3_2 | Peroxisome biogenesis protein 3-2 |
| 148 | MOS4_2 | Pre-mRNA-splicing factor SPF27 homolog |
| 148 | 27660 | hypothetical protein |
| 148 | GSTU4 | Glutathione S-transferase U4 |
| 148 | TAS_3 | Protein tas |
| 200 | 29900 | hypothetical protein |
| 200 | 29899 | hypothetical protein |
| 200 | SCD1_6 | DENN domain and WD repeat-containing protein SCD1 |
| 200 | 29897 | hypothetical protein |
| 200 | SCD1_6 | DENN domain and WD repeat-containing protein SCD1 |
| 200 | 29895 | hypothetical protein |
| 200 | ATK4_3 | Kinesin-4 |
| 200 | KP1_4 | Kinesin KP1 |
| 200 | ESP3_8 | Pre-mRNA-splicing factor ATP-dependent RNA helicase DEAH1 |
| 200 | ESP3_9 | Pre-mRNA-splicing factor ATP-dependent RNA helicase DEAH1 |
| 200 | ARP4_3 | Actin-related protein 4 |
| 200 | ARP4_4 | Actin-related protein 4 |
| 200 | Y39A1A | Ribosomal RNA small subunit methyltransferase nep-1 |
| 200 | 29883 | hypothetical protein |
| 305 | PDF2 | Defensin-like protein 1 |
| 305 | PDF2 | Defensin-like protein 6 |
| 305 | N_98 | TMV resistance protein N |
| 305 | N_84 | TMV resistance protein N |
| 305 | PDR2_6 | Probable manganese-transporting ATPase PDR2 |
| 305 | N_19 | TMV resistance protein N |
| 322 | UBP12_7 | Ubiquitin carboxyl-terminal hydrolase 12 |
| 322 | DLO2_6 | Protein DMR6-LIKE OXYGENASE 2 |
| 322 | AT5g09450_5 | Pentatricopeptide repeat-containing protein At5g09450, mitochondrial |
| 322 | AT5g09450_3 | Pentatricopeptide repeat-containing protein At5g09450, mitochondrial |
| 322 | DLO2_4 | Protein DMR6-LIKE OXYGENASE 2 |
| 336 | AT1g64760_3 | Glucan endo-1,3-beta-glucosidase 8 |
| 343 | UBP12_3 | Ubiquitin carboxyl-terminal hydrolase 12 |
| 343 | DLO2AT5g09450 | Protein DMR6-LIKE OXYGENASE 2 |
| 384 | 33334 | hypothetical protein |
| 384 | N54 | TMV resistance protein N |
| 384 | PDR2MYB308 | Probable manganese-transporting ATPase PDR2 |
| 384 | AT4g27220_44 | Probable disease resistance protein At4g27220 |
| 387 | MYB330 | Myb-related protein 330 |
| 388 | RGA4_58 | Putative disease resistance protein RGA4 |
| 388 | BMS1_10 | Ribosome biogenesis protein BMS1 homolog |
| 388 | RGA3_3 | Putative disease resistance protein RGA3 |

| | | |
|---|---|---|
| 388 | RGA4_14 | Putative disease resistance protein RGA4 |
| 388 | 33367 | hypothetical protein |
| 388 | BMS1_5 | Ribosome biogenesis protein BMS1 homolog |
| 464 | ACD6_15 | Protein ACCELERATED CELL DEATH 6 |
| 464 | 33929 | hypothetical protein |
| 493 | 34069 | hypothetical protein |
| 502 | ZFN1_2 | Zinc finger CCCH domain-containing protein ZFN-like |
| 502 | PIN5_3 | Auxin efflux carrier component 5 |
| 502 | CBSX6_2 | CBS domain-containing protein CBSX6 |
| 539 | AT2g19130b120 | G-type lectin S-receptor-like serine/threonine-protein kinase B120 |
| 590 | M3KE1_10 | MAP3K epsilon protein kinase 1 |
| 590 | M3KE1_36 | MAP3K epsilon protein kinase 1 |
| 590 | 34462 | hypothetical protein |
| 590 | CTSB_2 | Cathepsin B |
| 1389 | 35922 | hypothetical protein |
| 464 | ACD6_15 | Protein ACCELERATED CELL DEATH 6 |
| 464 | 33929 | hypothetical protein |
| 563 | BGAL9 | Beta-galactosidase 9 |
| 563 | BGAL9NEK6 | Beta-galactosidase 9 |
| 574 | AT1g07650_8 | Probable LRR receptor-like serine/threonine-protein kinase At1g07650 |
| 580 | 34425 | hypothetical protein |
| 580 | 34427 | hypothetical protein |
| 580 | RNP1_12 | Heterogeneous nuclear ribonucleoprotein 1 |
| 595 | AT3g47200_66 | UPF0481 protein At3g47200 |
| 595 | OS08g0118900 | Probable adenylate kinase 7, mitochondrial |
| 595 | 34484 | hypothetical protein |
| 595 | AT3g03770_5 | Probable inactive leucine-rich repeat receptor-like protein kinase At3g03770 |
| 609 | WAP_2 | WPP domain-associated protein |
| 609 | FKBP16-1_2 | Peptidyl-prolyl cis-trans isomerase FKBP16-1, chloroplastic |
| 709 | XPO1 | Protein EXPORTIN 1A |
| 769 | NFYB8_3 | Nuclear transcription factor Y subunit B-8 |
| 769 | 34929 | hypothetical protein |

**Supplemental Fig. S1.** Whole genome synteny between *Salix viminalis* assembly and *Populus trichocarpa*. Forward alignments are drawn in blue and reverse alignments are drawn in red.

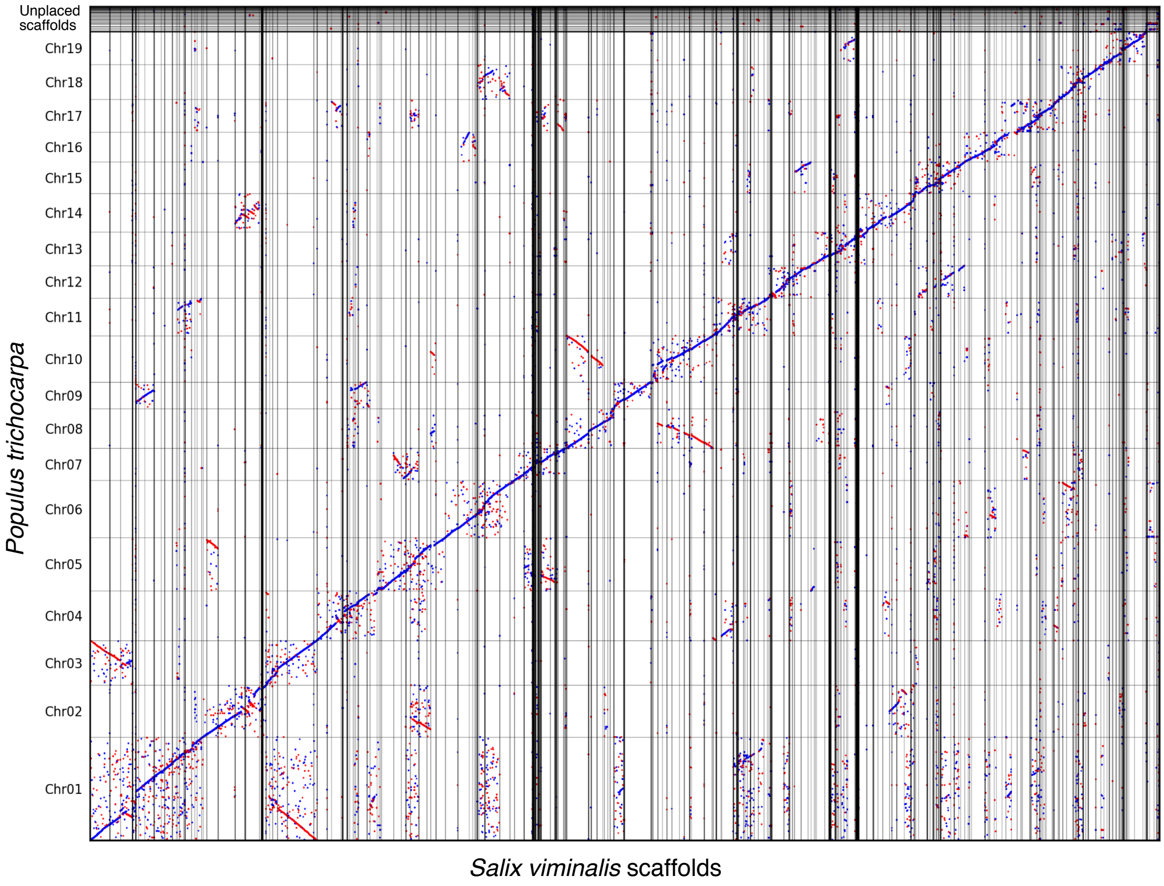

**Supplemental Fig S2**. Genetic markers aligned to chromosome 15 (from Pucholt et al. 2015) on our assembly. Markers associated with female segregation are filled in black.

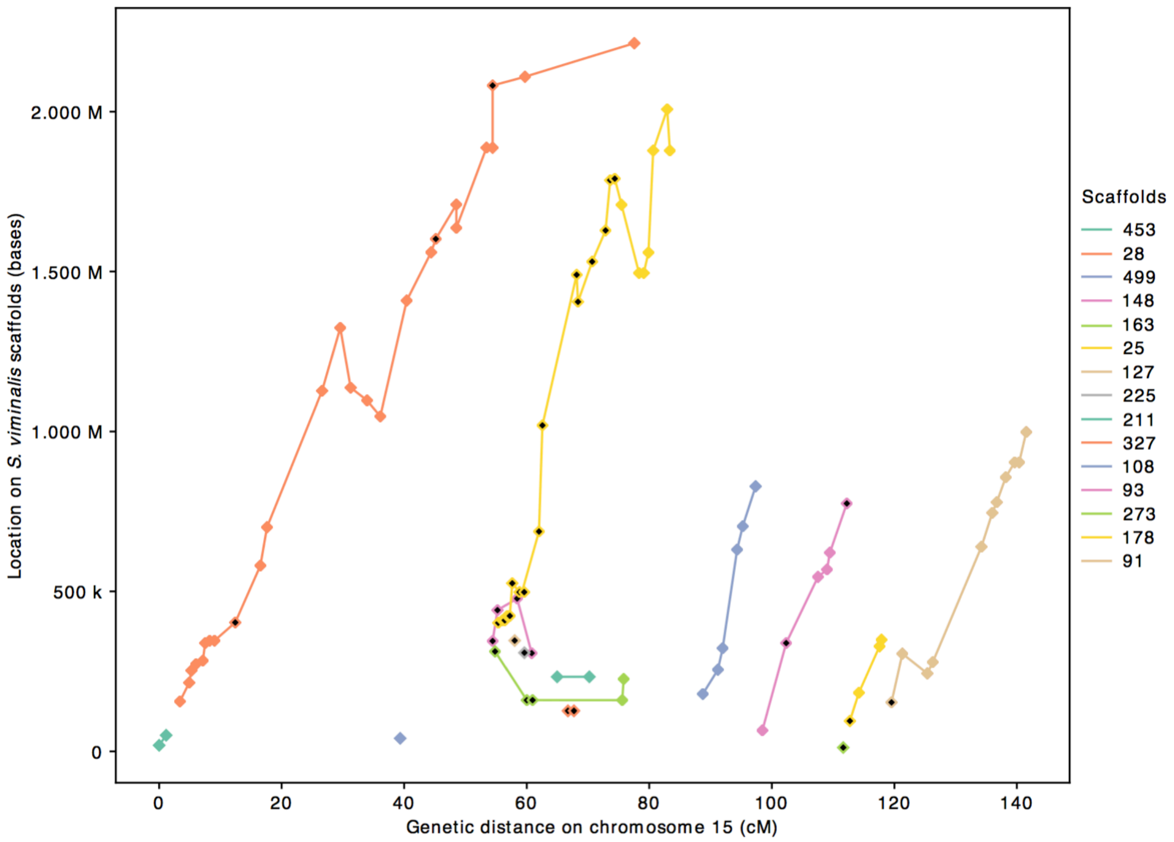

**Supplemental Fig S3**. Alignment between *Salix viminalis* and *Salix purpurea* SDR regions. One-to-one orthologous alignments between the *S. viminalis* scaffolds from chromosome 15 and the chromosome 15 of *S. purpurea*, with forward alignments drawn in blue and reverse alignments drawn in red. The SDR region of *S. purpurea* is delimited by the grey shaded area (10.7 Mb – 15.3 Mb, from Zhou et al. 2018). *S. viminalis* scaffolds anchored to chromosome 15 of *Populus trichocarpa* are highlighted in bold and those inferred to be part of the *S. viminalis* SDR are underlined.

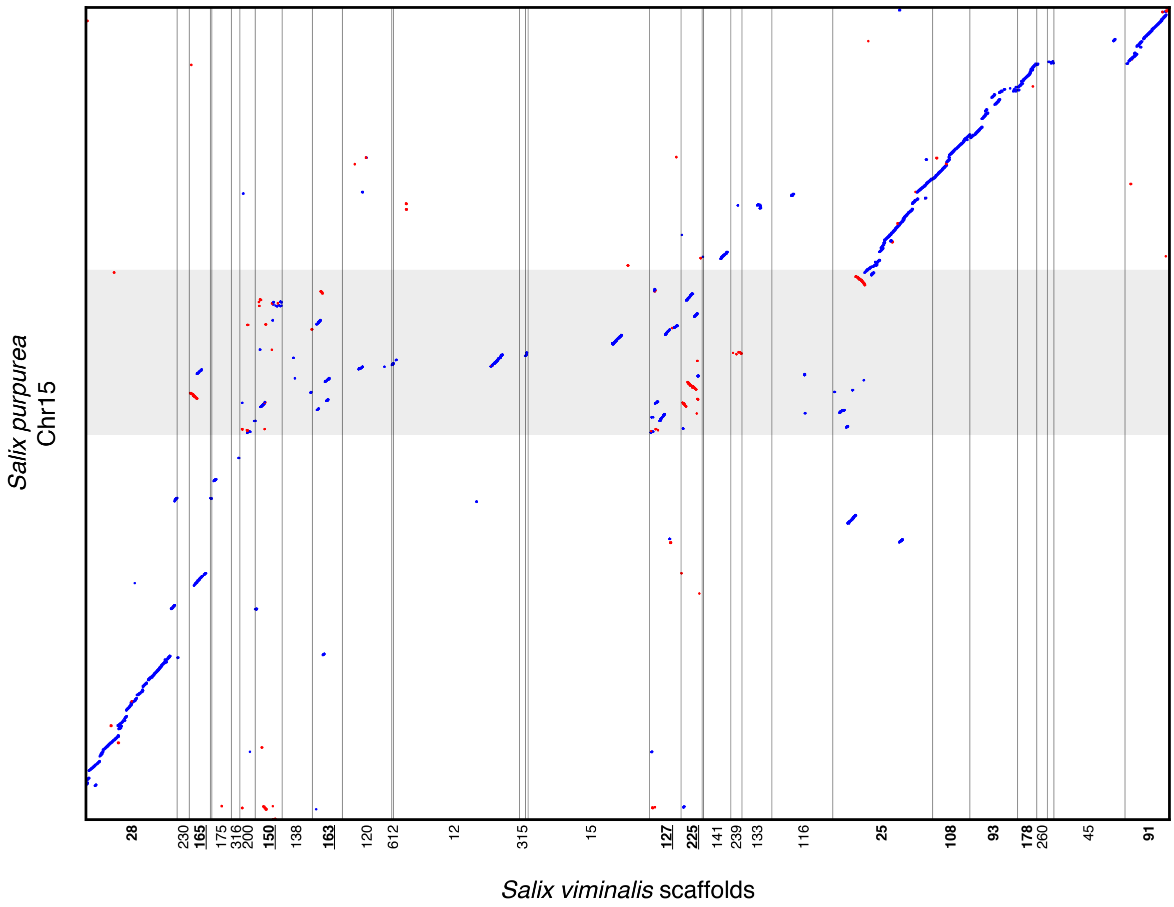

**Supplemental Fig. S4**. Percentage of fully phased haplotypes using 10X Genomic Chromium sequence data, across the whole genome (**A**) and chromosome 15 (**B**).

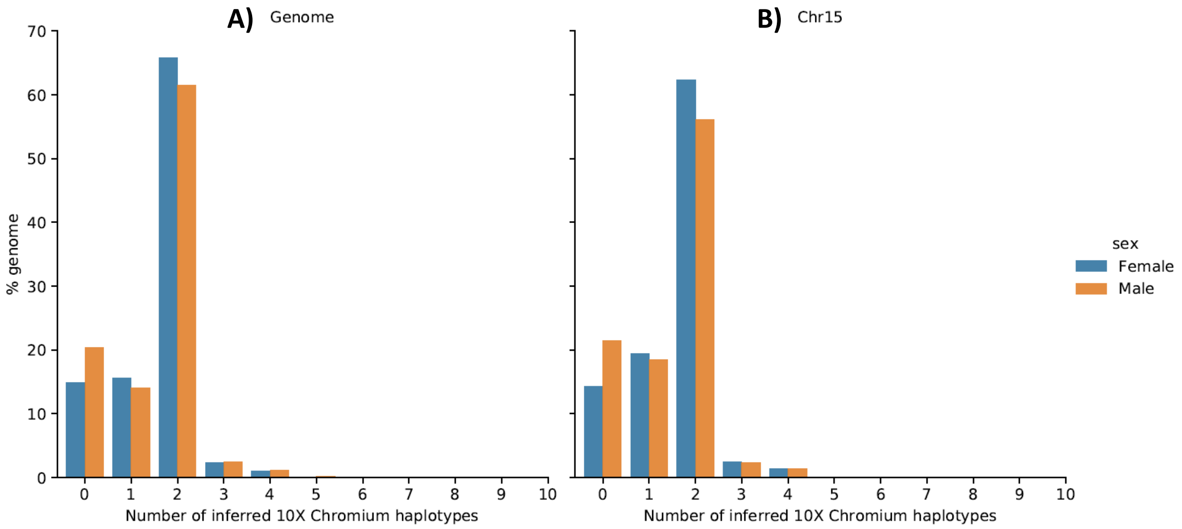

**Supplemental Fig. S5**. Phylogenetic trees between Z-W gene pairs in the basket willow SDR. Female *Salix viminalis* haplotypes are indicated with red squares and male haplotypes with blue squares. Trees were estimated by maximum likelihood. Bootstrap values >75% are indicated with black dots on the respective nodes. The poplar (*Populus trichocarpa*) ortholog was used to root the trees. The name of each gene is indicated at the top of the tree and the scaffold where the gene is located is indicated in parenthesis.

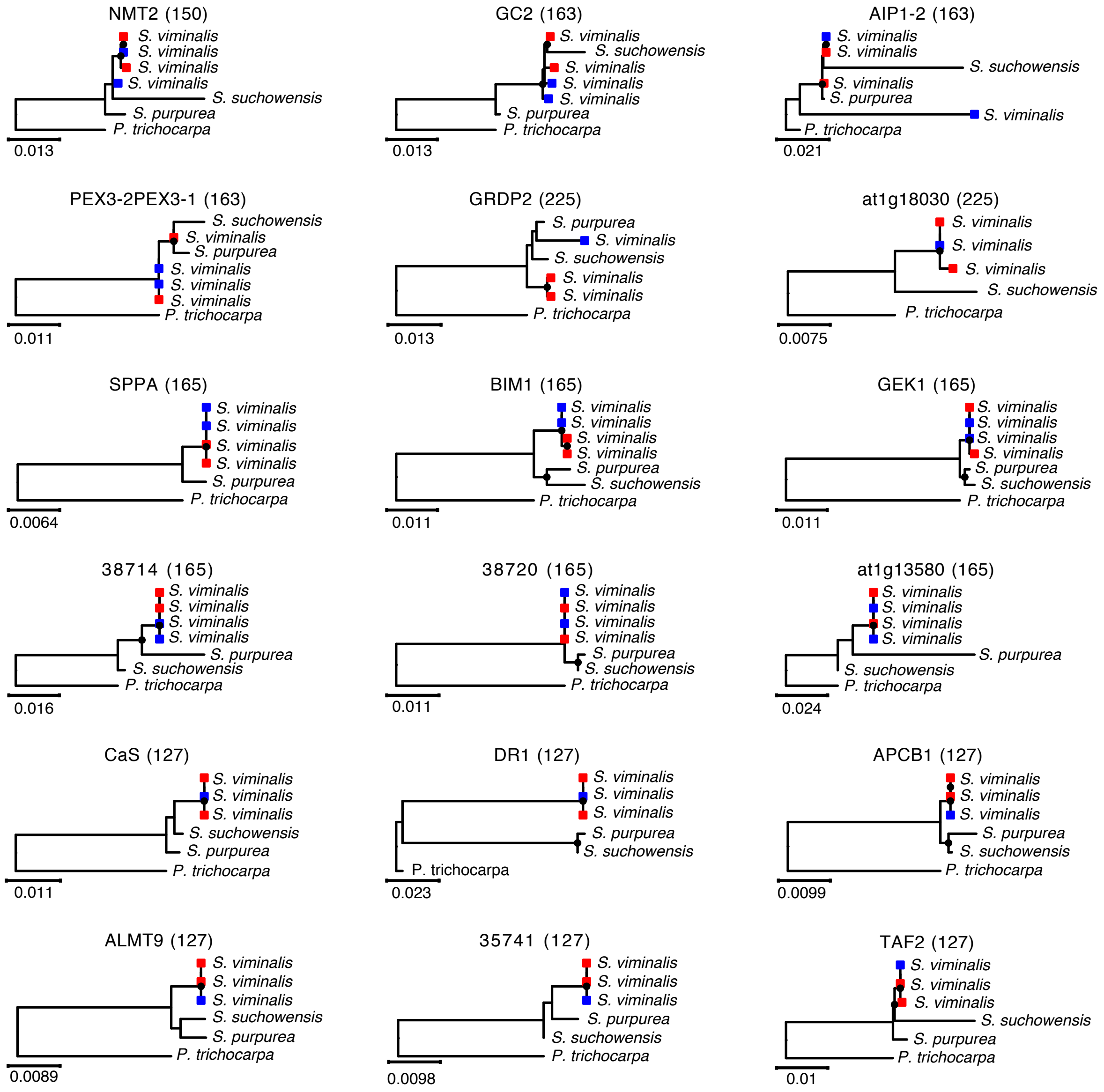

**Supplemental Fig. 6**. Density of repetitive elements across different genomic regions.

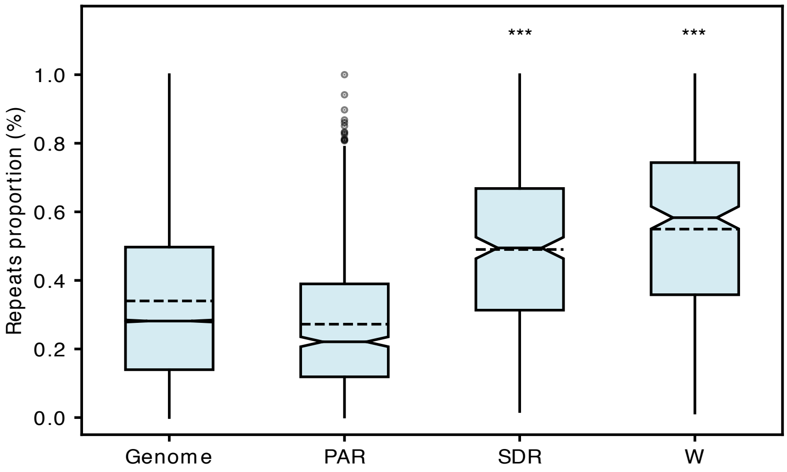
